# Orchestrating Self-Replication in Artificial Cells through Digital Microfluidics

**DOI:** 10.1101/2025.05.23.655734

**Authors:** Guanzhong Zhai, Pantelitsa Dimitriou, Jason Sengel, Mark Ian Wallace

## Abstract

A defining feature of living cells is their ability to self-replicate; but creating artificial cells with this capability remains challenging, due to the complexity of biological division machinery. Rather than seeking to reconstitute this machinery, here we take direct control of DNA replication and compartment division using digital microfluidics. This approach allows us to precisely orchestrate these two fundamental processes, providing insight into how they must be coupled for successful self-replication. Our system achieves autonomous cycles of replication and division, with daughter compartments inheriting parental DNA and maintaining genetic continuity across multiple generations - a key feature of living systems that has been difficult to achieve in artificial cells. By implementing these processes through direct physical manipulation rather than biochemical complexity, we provide a simple testbed that will help to disentangle the essential requirements for self-replicating systems.

## 1 Introduction

Artificial cells represent a major bottom-up approach to synthetic biology, with the goal of understanding living systems and mimicking their characteristics [1–3]. Recent advances have demonstrated artificial cells that exhibit key life-like characteristics, including genetic information transfer [4], metabolism [5], compartmentalisation [3], cell growth and division [6], and communication and motility [7]. Despite recent advancements, integrating all of these complex functions and establishing a continuous cell cycle of artificial cell replication remains a significant challenge [8]. Current approaches to compartmentalise DNA replication generally rely on biochemical mechanisms including protein-mediated membrane remodeling [9], phase separation [10], or chemical reactions that generate membrane components [11]. While these approaches have provided remarkable insights, they are frequently associated with low efficiency, poor control, and difficulty in achieving reliable coordination with DNA replication. To make progress, we reasoned that a system that enables explicit control over both replication and division would allow us to better understand the essential coupling between these processes, a fundamental question in understanding the minimal requirements for life-like systems. Thus here we sought to establish a digital microfluidic approach to directly control compartmentalisation and produce simple cell cycles capable of propagating genetic information.

Vesicle-based artificial cells have achieved DNA self-replication by encapsulating the appropriate constituents and exposing them to the required amplification temperatures [12–14]. Integrating gene replication with membrane dynamics can also enable the propagation of genetic information and the formation of spontaneous compartments [11]. However, vesicle division often requires extrinsic triggers, such as compositional changes of vesicle precursors [15], environmental modifications (e.g., illumination in the presence of photosensitive components [16]), osmotic pressure changes [17] to enhance vesicle fission, or pH shifts to drive vesicle growth [18, 19]. Notably, Kurihara et al. demonstrated self-assembly and DNA replication in giant unilamellar vesicles, where growth and division were driven by lipid precursor self-assembly [20]; although DNA replication over several cycles was reported, limitations were a lack of timing and size control during division. Abil et al. also recently encapsulated a DNA self-replicator in liposomes, where the DNA template could sustain self-encoded replication over *>* 10 rounds and carry out adaptive evolution through a recombinant gene expression system [21]; freeze-thaw cycles were used to achieve vesicle fusion and fission that requires redistribution of DNA across the vesicle population between evolutionary rounds. Artificial cells based on coacervates have also shown spontaneous division and growth [22], but predominantly without controlled inheritance of genetic information. For example, Dreher et al. demonstrated DNA-containing coacervates that could undergo division while maintaining DNA content but without control over the timing and coordination of these processes [23]. Coacervate droplets formed by in situ oligonucleotide ligation and elongation [24] suitable for Darwinian evolution [25] have also been reported, and more recently protocell growth induced by a compartmentalised DNA replication reaction [26].

DNA amplification within water-in-oil droplets have also been reported [27, 28]; For example, Sakatani et al. developed a simplified DNA replication system combining rolling-circle isothermal replication of DNA encoding a DNA polymerase (and externally supplied recombinase), in the presence of in vitro transcription and translation reagents to generate the encoded polymerase [29]. Thus although DNA self-replication in artificial cells are arguably well established, coordinating compartment division remains a challenge. For example, liposomes require some form of transmembrane channel to realise transport [30] and are often unable to maintain the encapsulate for prolonged periods [31, 32]. Coacervate’s advantage in lacking a phospholipid membrane also leads to restrictions in division due to surface tension requirements [33]. Similarly, emulsion systems are also membraneless, but have limited stability and size control in the absence of appropriate surfactant concentrations [34] and lack control of division and growth [27].

Overall, each of these methods possesses contrasting challenges that have hampered progress toward the reliable replication of genetic material and cell volume require to build a bottom-up minimal self-replicating cell. We reasoned that Digital microfluidics (DMF) provides an alternative route to directly control the size and shape of individual aqueous droplets and might serve as an alternative basis for artificial cell compartments [35, 36]. In DMF, the electrowetting-on-dielectric (EWOD) e”ect changes surface wettability resulting in controlled actuation of droplet position [37, 38]. DMF has low sample volume, digital control and flexible programmability, making it suitable for a wide range of biological applications - including single-cell proteomic [39], cell invasion [40], and cell culture [41]. Notably, DMF has revolutionised nucleic acid research, finding applications in DNA storage [42], RNA extraction [43], and DNA amplification [44]. Relevant to this work, Ruan et al. demonstrated amplification of DNA isolated from single cells within DMF droplets [45], and Kalsi et al. used DMF-based DNA amplification for gene detection [46].

Here, we adapt the open-source DMF platform OpenDrop [47] to generate autonomous cycles of DNA replication and compartment division (Fig. 1b). Open-Drop has been used to study areas as varied as the self-assembly of multifunctional magnetic nanoclusters [48], in situ spectroscopy [49], and cell-free protein expression and plasmid screening [50].

**Figure 1:**
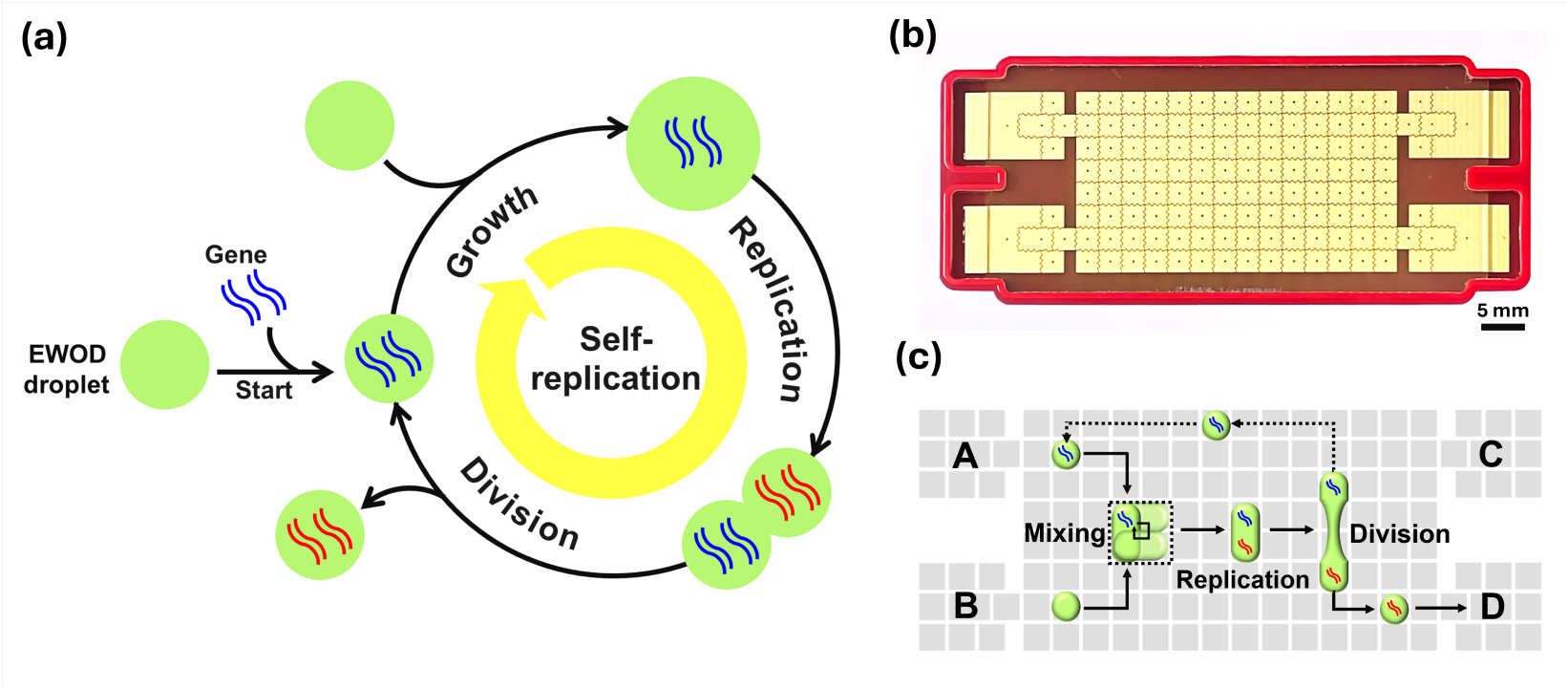
DMF-based DNA self-replication. (a) Schematic representation of the steps for DNA self-replication within EWOD droplets: DNA encapsulation, droplet growth, DNA replication within the droplet, and droplet division. (b) Photograph of the OpenDrop chip. Scale bar: 5 mm. OpenDrop comprises a rectangular array of 112 gold-coated electrodes, plus 4 input/output EWOD reservoirs. (c) Schematic diagram illustrating the replication process: Droplets containing reagents are combined and mixed to generate the droplet growth step. DNA replication is then achieved under temperature controlled conditions using resistive heater build into the array. The droplet is then divided equally. One daughter droplet is collected, while the other returns to the starting point to initiate the next round of self-replication.

Fig. 1a illustrates our overall strategy, in which genetic material is encapsulated inside an EWOD droplet and replication is initiated via programmed fusion with a second droplet containing the necessary components for DNA replication. The sequence of mixing, DNA replication, droplet division, and fusion then create a cycle of continuous self-replication (Fig. 1c). Through engineering of surface properties and temperature control, we have optimized the reliable DNA amplification within droplet-based cell mimics that can be precisely divided using EWOD actuation. Daughter droplets inherit the parental DNA and maintain replication capacity, enabling sustained multiple generations of self-replication. This system demonstrates a path toward artificial cells that can grow and multiply through direct physical control rather than biochemical complexity, providing insights into the minimal physical principles required for sustained self-replication.

## Results and Discussion

### DMF droplet replicator design

We first tackled compartment division by establishing a minimal cycle of droplet volume replication in the absence of DNA (Fig. 2). Replication cycling requires a programmed sequence of voltages to the electrodes in our array control the droplet position. Fig. 2a illustrates our final programmed sequence plus corresponding live camera images. Our minimal replication cycle comprised five stages: (i) **Initiation**: Reservoirs A and B each input a 2 µl droplet; (ii) **Mixing**: Droplets from A and B reservoirs were merged and repeatedly mixed (20 rounds clockwise mixing of a 2 *×* 2 square array; (iii) **Division**: Droplets were then split using a division array pattern (Fig. S5a) to create two equal sized daughters; (iv) **Growth**: One daughter droplet is returned to the mixing step for a subsequent round of fusion with a ‘fresh’ droplet from reservoir A. The second daughter droplet was collected and placed in a serpentine output queue where each droplet was spaced by three electrodes to avoid merging; (v) **Readout**: Finally the queue is read out, with individual droplets collected from the output reservoir using a gel loading pipette for further analysis.

**Figure 2:**
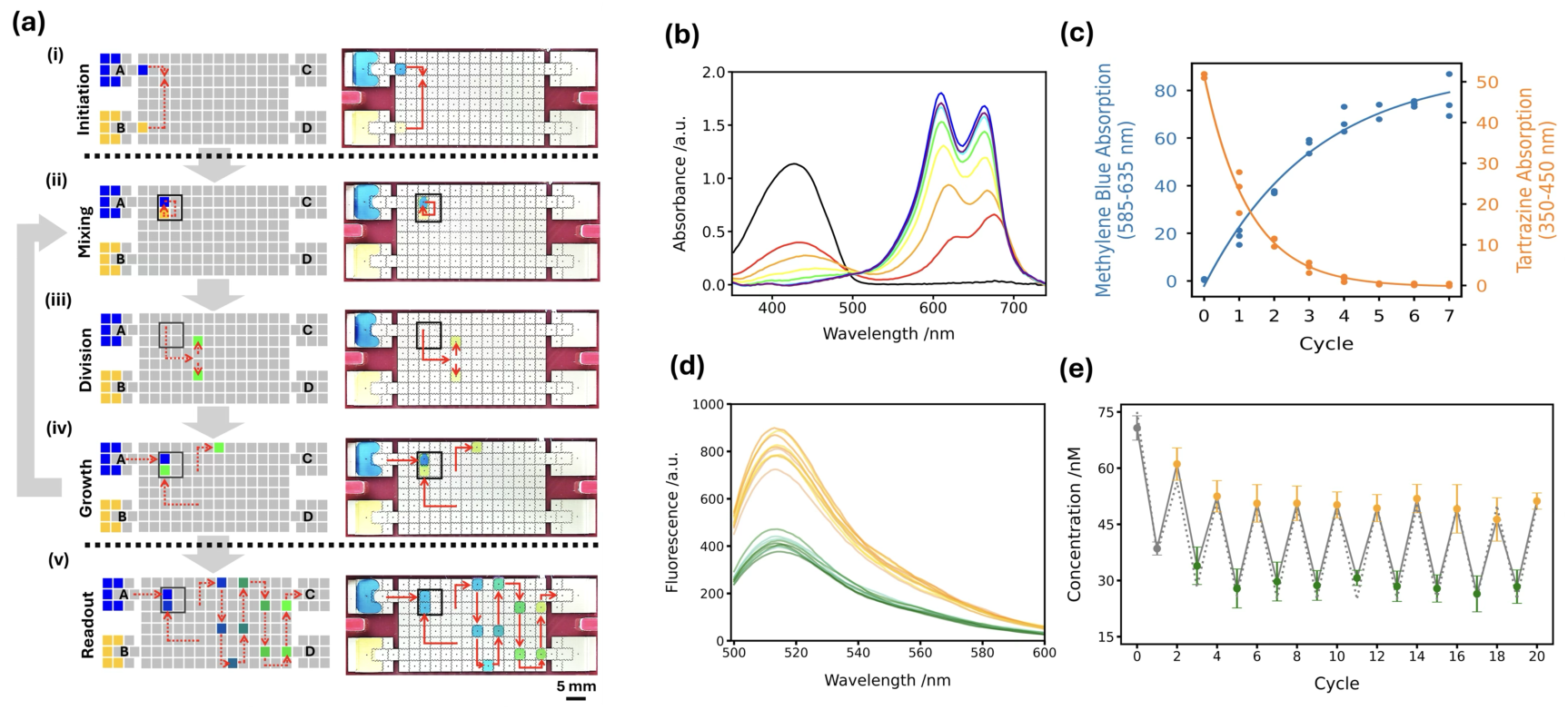
DMF cyclic replication. (a) Schematic (left) of the predefined electrode paths designed to generate droplet replication cycles. Red dashed lines indicate the paths followed by the droplets. Corresponding live camera images (right) are shown, where reservoirs A and B are filled with methylene blue and tartrazine, respectively. Scale bar: 5 mm. Droplet replication has five stages: (i) Initiation: Reservoirs A and B each input a 2 µl droplet. (ii) Mixing: Two droplets from the A and B reservoirs merged and repeatedly mixed for 20 rounds clockwise in a 2 *×* 2 electrode array. (iii) Division: Droplets are split into two via an electrode pattern designed to induce efficient division. (iv) Growth: One daughter droplet is returned to the Mixing stage and combined with a fresh droplet from reservoir A. The second daughter droplet is collected in a queue and stored for collection. Queue spacing is designed to avoid merging. (v) Readout: Droplets are sequentially output to reservoir C and collected using a gel loading pipette. (b) Rounds of cyclic droplet propagation by mixing methylene blue (reservoir A) with tartrazine (reservoir B). The resultant output droplets were analysed and the absorption spectra from serial dilution cycles are represented by a rainbow color gradient (from red (*×*1) to purple (*×*7)). Absorbance peaks at 427 and 664 nm correspond to tartrazine and methylene blue, respectively. The black line represents cycle 0, where the tartrazine droplet was not diluted. (c) Plotted relation between absorbance of tartrazine (orange) and methylene blue (blue) during droplet serial dilution cycles. (d) Spectra from two-state droplet cyclic replication (cycles 2–20) alternating between fluorescein (75 nM in 0.1 M Tris-HCl, pH 8.0, 1% (v/v) silicone oil AR5) and bu”er. Even-numbered cycles (fluorescein, orange) and odd-numbered cycles (bu”er, green) exhibit spectral overlap. (e) Two-state replication over 20 cycles. Measured (grey, solid line) and calculated (grey, dotted line) concentrations of fluorescein plotted per cycle. Colored markers show mean concentration over 3 repeats: cycles 0 and 1 (grey), even-numbered cycles (orange), odd-numbered cycles (green). Error bars: standard deviation.

### DMF replicator validation by serial mixing

To illustrate DMF replicator action rounds of cyclic droplet propagation were conducted by mixing methylene blue (1 mM, ultra-pure water, 1% v/v silicone oil AR5, reservoir A) with tartrazine (1 mM, ultra-pure water, 1% v/v silicone oil AR5, reservoir B) (Fig. 2a, Movie S1.) Two 2 µl droplets were generated - one from reservoir A (methylene blue) and one from reservoir B (tartrazine), and then mixed. For ‘cycle 0’, a droplet containing only trartrazine was generated. Output droplets from seven replication cycles were collected and absorption spectra determined (Fig. 2b). As expected, the characteristic absorption peak of tartrazine at 427 nm decreased and methylene blue at 664 nm increased as cycles repeated. During the replication cycles, methylene blue showed an exponential increase in absorbance, whereas tartrazine decreased exponentially and reached a plateau from cycle 5 onwards (Fig. 2c). The distinct trends of methylene blue and tartrazine are consistent with the expected serial mixing during the droplet replication cycles.

To further assess platform reproducibility, we also repeated cycles of mixing between an aqueous droplet of fluorescein (75 nM, 0.1 M Tris-HCl, pH 8.0, 1% (v/v) silicone oil AR5) and a droplet of bu”er alone (0.1 M Tris-HCl, pH 8.0, 1% (v/v) silicone oil AR5). Output droplet composition was then varied by alternating the introduced droplet between aqueous bu”er and the fluorescein reservoirs. The output droplets were extracted and the fluorescence spectra were measured (Fig. 2d). By calibrating measured fluorescence maxima against standardized fluorescein solutions (Fig. S6b) we can plot output droplet concentration over 20 cycles (Fig. 2e). Measured (grey solid line) and calculated (grey dashed line) concentrations after each programmed cycle are depicted. Fig. S7 shows the linear dependence between the measured and expected values. Across 5 independent repeats we measure a coefficient of variation of 3.72 % (27.39 ± 1.02 nM) (Fig. S8b) and 4.65 % (15.68 ± 0.73 nM) (Fig. S8d), demonstrating good reproducibility.

### DMF DNA replication

To develop a self-sustaining cycle of DNA replication, we next established conditions for e”ective DMF replication of DNA using recombinase polymerase amplification (RPA). We selected RPA as it operates at mild temperatures (37-42 °C), requires minimal sample preparation, and delivers rapid amplification [51–53]. In the RPA reaction mechanism, a recombinase ATPase (RecA) forms complexes with primers that facilitate their binding to complementary sequences on the target gene. After binding, RecA dissociates allowing DNA polymerase to initiate DNA synthesis; simultaneously, single-stranded binding protein (SSB) stabilizes the exposed DNA strands to prevent re-annealing [54]. The low operating temperature of RPA minimizes sample evaporation, making it particularly suitable for the low sample volumes present in microfluidic applications [46, 55]. We used a commercial fluorescent reporter (TwistAmp Exo, TwistDx) in which fluorescent signals are generated using a fluorophore/quencher ‘probe’ that is complementary to the target DNA [54, 56]. The probe also contains a tetrahydrofuran site that is a substrate for Exonuclease III - cutting the probe when complexed with target DNA separates the fluorophore/quencher complex and generates a fluorescent signal [46].

As RPA requires temperature control, we optimized OpenDrop for rapid DMF actuation of aqueous droplets not in air, but in an oil bulk phase - where droplet evaporation is prevented. (Supplementary Methods S2.2, Fig. S9). Whilst silicone oil (5 cSt) is more commonly used in DMF systems to prevent evaporation [28, 36], here we tested a range of potential compounds, and found octamethyltrisiloxane to provide superior droplet mobility and division into daughter droplets; as with a viscosity of 1 cSt, octamethyltrisiloxane is much closer to the kinematic viscosity of water (0.89 cSt, [57]). To achieve efficient droplet actuation, we also made a number of additional improvements: engineering a spacer to maintain a consistent distance between the electrode array and the ITO coverglass, and optimizing the thickness and composition of the polymer film coatings (Supplementary Methods S2.2, Fig. S1-S3).

Droplets were dispensed from reservoir A (RPA reaction components: 0.84 µM forward primer, 0.84 µM reverse primer, 0.24 µM probe, 0.32 nM positive control DNA template; 50 mM Tris-HCl, pH 7.9; 200 µM dNTPs; 3 mM ATP; 100 mM potassium acetate; 2 mM DTT; 50 mM phosphocreatine; 0.1% Tween 20 (v/v); a lyophilized enzyme pellet containing recombinase, SSB , strand-displacing polymerase, and ExoIII) and reservoir B (28 mM magnesium acetate; 50 mM Tris-HCl, pH 7.9; 200 µM dNTPs; 3 mM ATP; 100 mM potassium acetate; 2 mM DTT; 50 mM phosphocreatine; 0.1% Tween 20 (v/v)). Droplets were merged and mixed (Fig. 3a), initiating RPA. The mixed droplet was moved to a resistive heating element and maintained at 40 °C (Fig. 3a). RPA was monitored by collecting the output droplets at 10-minute intervals and measuring the fluorescence response of the probe (Fig. 3b). To monitor amplification kinetics, a series of independent DMF RPA reactions were performed for time points: 0, 10, 20, 30, 40, 50, and 60 minutes. For each time point, fresh reaction droplets were dispensed from reservoirs A and B, respectively, merged and mixed to initiate amplification, and then incubated at 40 °C. The amplified products were collected and analyzed. Fig. 3c shows that the reaction increases, reaching a plateau ( 50 mins) as reagents are consumed. We also repeated the DMF reaction without exonuclease to allow gel electrophoresis of the DNA products [58, 59]. Here, A 277 bp segment of pUC19, was amplified as previously described [60]. Agarose gel electrophoresis of on-chip RPA products confirms a time-dependent amplification of DNA (Fig. 3d).

**Figure 3:**
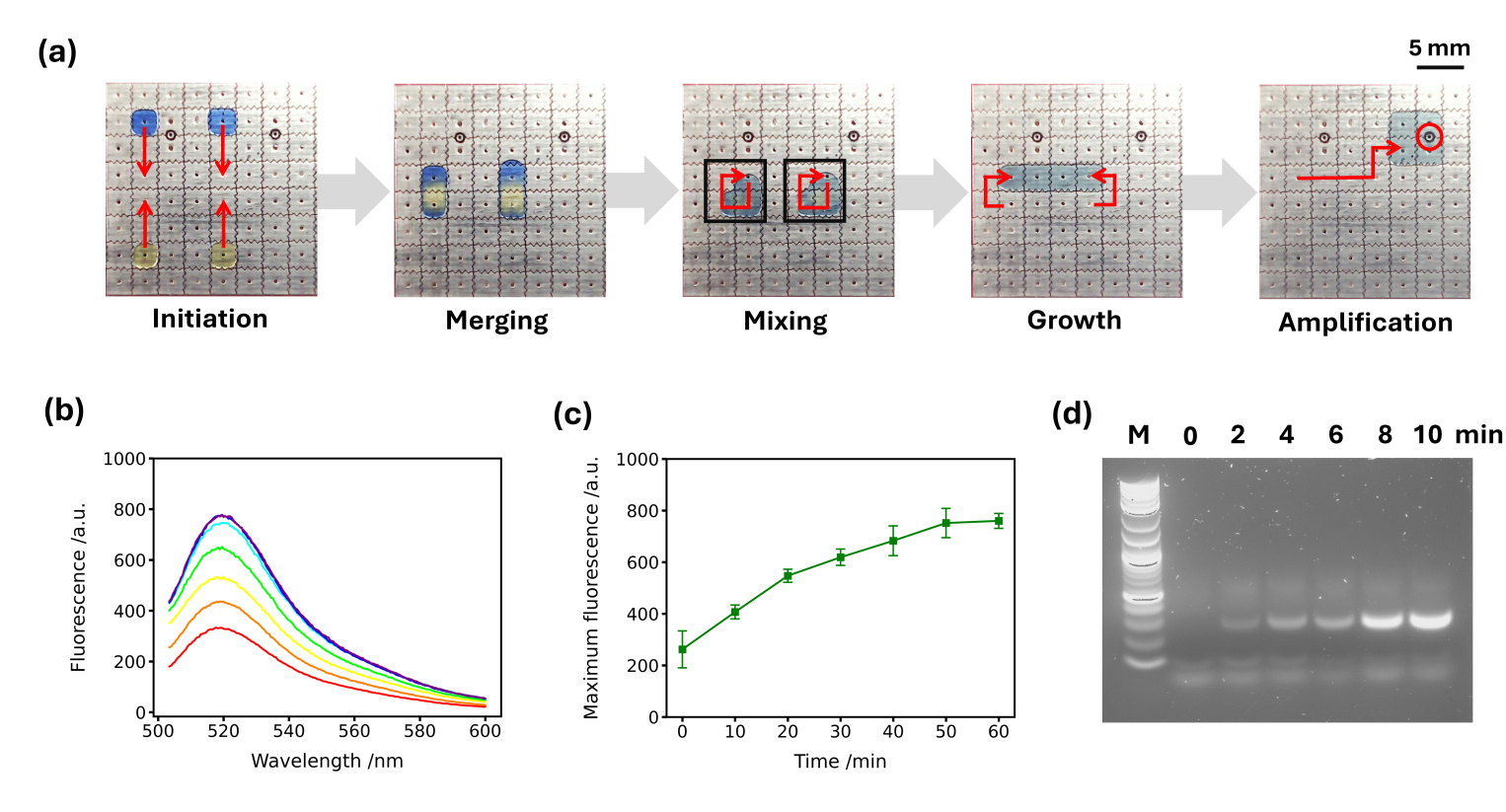
DMF RPA amplification. (a) Schematic of the steps required for on-chip RPA. Methylene blue and tartrazine (yellow) have been added to the droplets in these images to improve visibility. Initiation: Two (blue) droplets are created from the RPA mixture and two (yellow) droplets from the Mg(OAc)_2_ solution. Merging and Mixing: Blue and yellow droplets were merged, and mixed within two square regions. Growth: Four green droplets merged together after the completion of mixing. Amplification: Droplets were heated to 40 °C on the chip heater (red circle). (b) Representative fluorescence spectra of RPA reaction on a chip. 0 min (red), 10 min (orange), 20 min (yellow), 30 min (green), 40 min (cyan), 50 min (blue), 60 min (purple). (c) Kinetics of RPA reaction, reporting fluorescence maxima every 10 minutes. The error bars represent the standard deviation, *n* = 3. (d) Agarose gel electrophoresis of RPA kinetic products from pUC19.

### Sustained DMF DNA replication in an artificial cell cycle

Finally, we combined these approaches to determine conditions for sustained cycles of replication and division using DMF. We first quantified how Mg^2+^ and EDTA activate and deactivate DMF RPA (Supplementary Methods S2.3). Fig. 4a illustrates the DMF program. To initiate the DMF program, in cycle 0 an 8 µL droplet from the DNA solution (Reservoir C) was mixed with a 6 µL droplet of RPA reaction mixture and a 6 µL droplet of Mg(OAc)_2_ solution (Fig. 4a,(i)). The volume was then divided into two equal daughters, one of which was collected for analysis, the second entered cycle 1. In cycle 1, the cycle 0 daughter droplet was again combined with 4 µL from the RPA reaction mixture (Reservoir A), 6 µL of the Mg(OAc)_2_ solution (Reservoir B). The combined reaction volume was mixed, before moving to a resistive heating element and heated on chip at 40 °C for 15 minutes (Fig. 4a,(ii)). A total of 20 cycles were repeated, where the concentrations and volume of the RPA reaction mixture added in each cycle remained constant (to ensure the same supply of primers, probe, enzymes, ATP, and other components). 6 µL of Mg(OAc)_2_ solution (56 mM magnesium acetate, 67.76 mM Tris-HCl, 271.04 µM dNTPs, 4.07 mM ATP, 135.52 mM potassium acetate, 2.71 mM DTT, 67.76 mM phosphocreatine, and 0.1% Tween 20 (v/v), pH 7.9) was added in cycles 1-5 and 11-15, and 6 µL of EDTA solution (160 mM EDTA, 67.76 mM Tris-HCl, 271.04 µM dNTPs, 4.07 mM ATP, 135.52 mM potassium acetate, 2.71 mM DTT, 67.76 mM phosphocreatine, and 0.1% Tween 20 (v/v), pH 7.9) was supplied in cycles 6-10 and 16-20.

**Figure 4:**
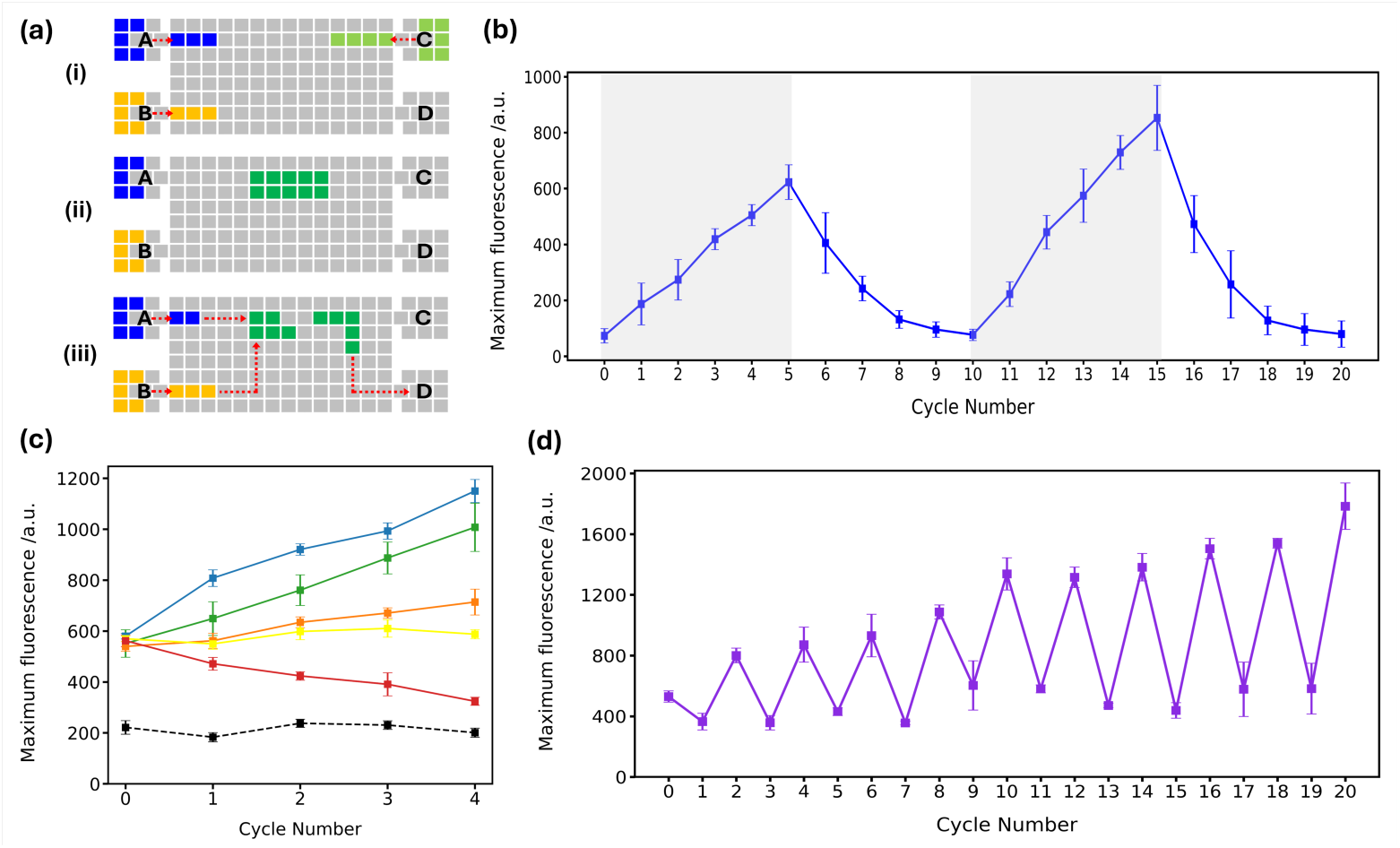
Regulation of the DNA self-replication. (a) Schematic of DMF program for DNA self-replication: (i) 6 µL droplets were dispensed from Reservoirs A (RPA reaction mixture) and B (Mg(OAc)_2_ solution), respectively. Reservoir C input 8 µL droplet of the DNA template only in cycle 0. (ii) The 20 µL mixture was mixed and transported to a heating element for incubation (no heating in cycle 0). (iii) The reaction droplet was divided into two equal 10 µL daughter droplets. One was collected via Reservoir D, and the other entered the next cycle. The remaining daughter droplet was mixed with fresh inputs: 4 µL from Reservoir A and 6 µL from Reservoir B, mixed and heated at 40 °C. (b) Periodic addition of Mg(OAc)_2_ (Reservoir B, cycles 1-5 and 11-15) and EDTA (Reservoir B, cycles 6-10 and 16-20) enabled regulated amplification and attenuation of DNA. Fluorescence increased during Mg(OAc)_2_ addition (grey area) and decreased during EDTA addition due to inhibited amplification. (c) Fluorescence intensity corresponding DMF self-replication cycles at di”erent temperatures. 25 °C (red), 30 °C (yellow), 35 °C (orange), 45 °C (green), and 55 °C (blue). A baseline control, consisting of the reaction mixture without a DNA template, was maintained at the optimal temperature of 40 °C (black dashed line). (d) Maximum fluorescence intensity during cycles of temperature switching. Cycle 0 represents the starting point without any amplification. For odd-numbered cycles (cycle 1, cycle 3,… cycle 19), reaction temperature was set to 25 °C. For even-numbered cycles (cycle 2, cycle 4,… cycle 20), the temperature was set to 45 °C. Each cycle supplies the same components and concentrations as in (a), stage (iii).

As shown in Fig. 4b, fluorescence signals exhibit an increasing trend during cycles 1-5 and 11-15, indicating DNA amplification. During cycles 6-10 and 16-20, fluorescence signals decreased significantly, showing that EDTA e”ectively chelated Mg^2+^, thus inhibiting DNA amplification, and reducing signals as DNA is serially diluted in each cycle. Each cycle demonstrates an integrated minimal cell cycle [8] achieved through DMF actuation.

With a minimal DMF cell cycle established, we sought to optimize conditions to enable continuous DNA propagation over multiple cycles. As RPA is highly temperature dependent [54], we examined DNA self-replication cycles at di”erent temperatures as a means to control reaction propagation. To achieve this, we used the DMF program described in Fig. 4a. Fig. 4c shows the temperature dependence of fluorescence intensity; for the reagent concentrations and droplet volumes optimized in this example, at approximately 40 °C the fluorescence intensity minimally increased, indicating a balance between RPA and the volume reduction upon cell division - leaving the concentration of DNA in each cycle unchanged. Based on these results, we then performed temperature-switching experiments over 20 rounds: The experimental setup was identical to Fig. 4a, (i)-(iii). Temperatureswitching began from cycle 1, where the reaction temperature was set at 25 °C for odd-numbered cycles (cycle 1, cycle 3, etc.), while for even-numbered cycles (cycle 2, cycle 4, etc.), the temperature was maintained at 45 °C. Fig. 4d shows continuous modulation of replication efficiency over the 20 cycles of DMF droplet division and replication, enabling long-term maintenance of DNA concentration over continuous cycles of DMF-based artificial cell replication.

## Conclusion

This work establishes DMF as a powerful platform for creating artificial cells with controlled cycles of replication and division - two defining features of living systems. By decoupling the biochemical complexity of natural cells from their essential physical processes, our approach o”ers a framework to explore how genetic replication can be coordinated with compartment division for successful propagation of information. Our approach is also a suitable analogue for early self-replicating systems that may have relied more on physical principles than biochemical sophistication, before evolving the elaborate protein machinery seen in modern cells. Our platform enables control over experimental parameters relevant for investigating the emergence of evolutionary behaviors, including manipulation of selection pressures, inheritance patterns, and mutation rates; capabilities that are challenging to access in conventional artificial cell systems. Although here we have focused on robust, reproducible readout of DNA, we envisage an on-chip readout, such as fluorescence, would enable cyclic rounds of selected evolution using this approach. By providing a reliable, programmable testbed for studying requirements of self-replicating systems, this work opens new avenues for exploring the boundaries between non-living and living matter - providing an alterative route to circumvent many of the limitations of membrane-containing and membrane-less systems in studying the mechanics of replication relevant to artificial cells.

## Materials and Methods

### Materials

OpenDrop V4 driver board, drilled ITO coverglasss, and cartridge frames were purchased from GaudiLabs LLC (Switzerland). FluoroPel PFC1601V was purchased from CYTONIX (USA). Octamethyltrisiloxane was obtained from Fluorochem Limited (UK). Methylene Blue hydrate (*↑*95%), Tartrazine (*↑*85%), Tris(hydroxymethyl)aminomethane (*↑*99.8%), and silicone oil AR5 (5 cSt at 25 °C) were purchased from Sigma-Aldrich (UK/USA). Fluorescein (free acid) was obtained from Fluka Chemie GmbH (UK). TwistAmp exo and basic RPA kits were purchased from TwistDx (USA). pUC19 was obtained from New England BioLabs (UK).

### OpenDrop device setup and operation

The OpenDrop device comprises of a 14 *×* 8 array of gold-coated electrodes (2.75 mm *×* 2.75 mm, with 4 mil gaps) and integrated resistive heating elements. The platform was operated in alternating current (AC) mode at 160 V and 350 Hz for droplet manipulation experiments, and at 50 V and 100 Hz for heating operations. Each electrode accommodated a 2 µL droplet volume, with reservoirs holding 16 µL. Modifications to the standard device were made to enable e”ective DMF-based artificial cell replication as detailed in the Supplementary Information.

### Serial cyclic dilution setup and operation

Reservoir A and B were injected with 16 µL of methylene blue (1 mM, ultrapure water, 1% v/v silicone oil AR5) and 16 µL of tartrazine (1 mM, ultra-pure water, 1% v/v silicone oil AR5), respectively. The programmed pattern consisted of five steps: (i) Initiation, (ii) Mixing, (iii) Division, (iv) Growth and (v) Readout. The droplet for cycle 0 was a 1 mM tartrazine solution, and the droplets for cycles 1 to 7 were continuously diluted with 1 mM methylene blue solution.

For sequential droplet growth and division experiments, Reservoir A was filled with 16 µL of fluorescein (25 nM in 0.1 M Tris-HCl bu”er, pH 8.0) containing 1% (v/v) silicone oil AR5, and Reservoir B with 16 µL of 0.1 M Tris bu”er (pH 8.0) containing 1% (v/v) silicone oil AR5. The droplets were sequentially analyzed using a fluorometer (Cary Eclipse, excitation wavelength 490 nm, emission wavelength 513 nm, excitation slit 5 nm, emission slit 5 nm, PMT voltage 1000 V).

### Real-time RPA reaction on DMF setup and operation

RPA mixture was prepared in a 2 mL sterile microcentrifuge tube, consisting of 1.68 µL of 10 µM forward primer, 1.68 µL of 10 µM reverse primer, 0.48 µL of 10 µM probe, 5.6 µL of rehydration bu”er, 0.8 µL of 8.06 nM positive control template, 7.76 µL of autoclaved water, and 2 µL of 1% Tween 20 solution. The 20 µL RPA mixture was centrifuged at 7,000 rpm (3,200 *×* g) for 90 seconds, after which it was carefully introduced to a lyophilized enzyme pellet. This preparation was centrifuged at 6,000 rpm (2,000 *×* g) for 3 minutes and added to reservoir A. A diluted Mg(OAc)_2_ solution was prepared by mixing 2 µL of 280 mM Mg(OAc)_2_ with 18 µL of rehydration bu”er, centrifuged at 7,000 rpm (3,200*×* g) for 90 seconds and loaded into reservoir B. The chip was filled with 800 µL of octamethyltrisiloxane. Droplets were distributed from reservoirs A and B, mixed for 4 minutes, and heated to 40 °C. Samples were collected at specified time points, placed on ice to quench the reaction, and analyzed by fluorescence or gel electrophoresis.

For gel electrophoresis analysis, a 277 bp segment of pUC19 was amplified using a TwistAmp Basic kit with primers F41 (5’-GGGTAACGCC AGGGTTTTCC CAGTCACGAC GTTGTAAAAC G-3’) and R-43-mer (5’-ACAGGTTTCC CGACTGGAAA GCGGGCAGTG AGCGCAACGC-3’). Products were analyzed on a 1.5% (w/v) agarose gel containing SYBR Gold at 60 V for 90 minutes.

### Regulation of sustained DMF DNA self-replication

For Mg^2+^/EDTA switching experiments, a RPA mixture was loaded into reservoir A (containing primers, probe, rehydration bu”er, Tween 20 and lyophilized enzyme), while reservoir B contained either 56 mM Mg(OAc)_2_ or 160 mM EDTA solution. DNA template (0.32 nM) was added to reservoir C. In cycle 0, droplets of DNA solution were mixed with RPA mixture and Mg(OAc)_2_ without amplifi-cation, divided into two equal volumes, with one half collected for analysis and the other entering cycle 1. In subsequent cycles, Mg(OAc)_2_ was supplied in cycles 1-5 and 11-15, while EDTA was supplied in cycles 6-10 and 16-20. Each cycle was heated at 40 °C for 15 minutes.

For temperature-switching experiments, reaction temperature alternated between 25 °C for odd-numbered cycles and 45 °C for even-numbered cycles, with Mg(OAc)_2_ supplied in every cycle. The DMF platform operated at 160 V at 200 Hz for droplet manipulation and 50 V at 100 Hz for heating. All experiments were performed in triplicate, with samples analyzed using a fluorometer (Cary Eclipse, excitation wavelength 488 nm, emission wavelength 520 nm, excitation slit 10 nm, emission slit 10 nm, PMT voltage 900 V).

## Supporting information

Supplementary Methods and Results

## Acknowledgments

Zhai thanks the King’s-China Scholarship Council PhD Scholarship program for funding. Wallace and Dimitriou are supported by the Wellcome Trust (224327/ Z/21/Z).

## Author Contributions

Zhai designed and performed all experiments, analyzed the data, and drafted the manuscript. Dimitriou assisted with data analysis and manuscript preparation. Sengel established the original OpenDrop platform and provided technical support. Wallace conceived the project, secured funding, supervised the research, and finalized the manuscript. All authors reviewed and approved the final version.

## References

1. Tang TC, An B, Huang Y, et al. Materials design by synthetic biology. Nature Reviews Materials 2021;6:332–50.

2. Schwille P. Bottom-up synthetic biology: engineering in a tinkerer’s world. Science 2011;333:1252–4.

3. Schwille P, Spatz J, Landfester K, et al. MaxSynBio: avenues towards creating cells from the bottom up. Angewandte Chemie International Edition 2018;57:13382–92.

4. Samanta A, Baranda Pellejero L, Masukawa M, et al. DNA-empowered synthetic cells as minimalistic life forms. Nature Reviews Chemistry 2024;8:1–17.

5. Bailoni E, Partipilo M, Coenradij J, et al. Minimal out-of-equilibrium metabolism for synthetic cells: a membrane perspective. ACS Synthetic Biology 2023;12:922–46.

6. Jiang W, Wu Z, Gao Z, et al. Artificial cells: past, present and future. ACS nano 2022;16:15705–33.

7. Wang L, Song S, Hest J van, et al. Biomimicry of cellular motility and communication based on synthetic soft-architectures. Small 2020;16:1907680.

8. Olivi L, Berger M, Creyghton RN, et al. Towards a synthetic cell cycle. Nature Communications 2021;12:4531.

9. Sharma B, Moghimianavval H, Hwang SW, et al. Synthetic cell as a platform for understanding membrane-membrane interactions. Membranes 2021;11:912.

10. Matsuo M and Kurihara K. Proliferating coacervate droplets as the missing link between chemistry and biology in the origins of life. Nature Communications 2021;12:5487.

11. Kurihara K, Tamura M, Shohda Ki, et al. Self-reproduction of supramolecular giant vesicles combined with the amplification of encapsulated DNA. Nature Chemistry 2011;3:775–81.

12. Sato Y, Komiya K, Kawamata I, et al. Isothermal amplification of specific DNA molecules inside giant unilamellar vesicles. Chemical Communications 2019;55:9084–7.

13. Lee S, Koo H, Na JH, et al. DNA amplification in neutral liposomes for safe and e!cient gene delivery. ACS nano 2014;8:4257–67.

14. Van Nies P, Westerlaken I, Blanken D, et al. Self-replication of DNA by its encoded proteins in liposome-based synthetic cells. Nature Communications 2018;9:1583.

15. Matsuo M, Kan Y, Kurihara K, et al. DNA length-dependent division of a giant vesicle-based model protocell. Scientific Reports 2019;9:6916.

16. Dreher Y, Jahnke K, Schröter M, et al. Light-triggered cargo loading and division of DNA-containing giant unilamellar lipid vesicles. Nano Letters 2021;21:5952–7.

17. Liu X, Stenhammar J, Wennerström H, et al. Vesicles balance osmotic stress with bending energy that can be released to form daughter vesicles. The Journal of Physical Chemistry Letters 2022;13:498–507.

18. Suzuki K, Kurihara K, Okura Y, et al. pH-induced switchable vesicular aggregation of zwitterionic and anionic phospholipids. Chemistry Letters 2012;41:1084–6.

19. Suzuki K, Aboshi R, Kurihara K, et al. Adhesion and fusion of two kinds of phospholipid hybrid vesicles controlled by surface charges of vesicular membranes. Chemistry Letters 2012;41:789–91.

20. Kurihara K, Okura Y, Matsuo M, et al. A recursive vesicle-based model protocell with a primitive model cell cycle. Nature Communications 2015;6:8352.

21. Abil Z, Restrepo Sierra AM, Stan AR, et al. Darwinian evolution of self-replicating DNA in a synthetic protocell. Nature Communications 2024;15:9091.

22. Slootbeek AD, Haren MH van, Smokers IB, et al. Growth, replication and division enable evolution of coacervate protocells. Chemical Communications 2022;58:11183–200.

23. Dreher Y, Jahnke K, Bobkova E, et al. Division and regrowth of phaseseparated giant unilamellar vesicles. Angewandte Chemie International Edition 2021;60:10661–9.

24. Fraccia TP and Martin N. Non-enzymatic oligonucleotide ligation in coacervate protocells sustains compartment-content coupling. Nature Communications 2023;14:2606.

25. Adamski P, Eleveld M, Sood A, et al. From self-replication to replicator systems en route to de novo life. Nature Reviews Chemistry 2020;4:386–403.

26. Minagawa Y, Yabuta M, Su’etsugu M, et al. Self-growing protocell models in aqueous two-phase system induced by internal DNA replication reaction. Nature Communications 2025;16:1522.

27. Ueno H, Sawada H, Soga N, et al. Amplification of over 100 kbp DNA from single template molecules in femtoliter droplets. ACS Synthetic Biology 2021;10:2179–86.

28. Laos R and Benner S. Fluorinated oil-surfactant mixtures with the density of water: artificial cells for synthetic biology. Plos One 2022;17:e0252361.

29. Sakatani Y, Yomo T, and Ichihashi N. Self-replication of circular DNA by a self-encoded DNA polymerase through rolling-circle replication and recombination. Scientific Reports 2018;8:13089.

30. Lu Y, Allegri G, and Huskens J. Vesicle-based artificial cells: materials, construction methods and applications. Materials Horizons 2022;9:892–907.

31. Ha Y, Koo Y, Park SK, et al. Liposome leakage and increased cellular permeability induced by guanidine-based oligomers: e”ects of liposome composition on liposome leakage and human lung epithelial barrier permeability. RSC Advances 2021;11:32000–11.

32. Wang X, Du H, Wang Z, et al. Versatile phospholipid assemblies for functional synthetic cells and artificial tissues. Advanced Materials 2021;33:2002635.

33. Lin Z, Beneyton T, Baret JC, et al. Coacervate droplets for synthetic cells. Small Methods 2023;7:2300496.

34. Torre P, Keating CD, and Mansy SS. Multiphase water-in-oil emulsion droplets for cell-free transcription–translation. Langmuir 2014;30:5695–9.

35. Ng AH, Fobel R, Fobel C, et al. A digital microfluidic system for serological immunoassays in remote settings. Science Translational Medicine 2018;10:eaar6076.

36. Xu X, Cai L, Liang S, et al. Digital microfluidics for biological analysis and applications. Lab on a Chip 2023;23:1169–91.

37. Li J and Kim CJ. Current commercialization status of electrowetting-ondielectric (EWOD) digital microfluidics. Lab on a Chip 2020;20:1705–12.

38. Kim JH, Lee JH, Mirzaei A, et al. Electrowetting-on-dielectric characteristics of ZnO nanorods. Scientific Reports 2020;10:14194.

39. Yang Z, Jin K, Chen Y, et al. AM-DMF-SCP: integrated single-cell proteomics analysis on an active matrix digital microfluidic chip. JACS Au 2024;4:1811–23.

40. Lamanna J, Scott EY, Edwards HS, et al. Digital microfluidic isolation of single cells for-Omics. Nature Communications 2020;11:5632.

41. Srigunapalan S, Eydelnant IA, Simmons CA, et al. A digital microfluidic platform for primary cell culture and analysis. Lab on a Chip 2012;12:369–75.

42. Antkowiak PL, Koch J, Nguyen BH, et al. Integrating DNA encapsulates and digital microfluidics for automated data storage in DNA. Small 2022;18:2107381.

43. Jebrail MJ, Sinha A, Vellucci S, et al. World-to-digital-microfluidic interface enabling extraction and purification of RNA from human whole blood. Analytical Chemistry 2014;86:3856–62.

44. Wan L, Chen T, Gao J, et al. A digital microfluidic system for loop-mediated isothermal amplification and sequence specific pathogen detection. Scientific Reports 2017;7:14586.

45. Ruan Q, Ruan W, Lin X, et al. Digital-WGS: automated, highly e!cient whole-genome sequencing of single cells by digital microfluidics. Science Advances 2020;6:eabd6454.

46. Kalsi S, Valiadi M, Tsaloglou MN, et al. Rapid and sensitive detection of antibiotic resistance on a programmable digital microfluidic platform. Lab on a Chip 2015;15:3065–75.

47. Alistar M and Gaudenz U. OpenDrop: an integrated do-it-yourself platform for personal use of biochips. Bioengineering 2017;4:45.

48. Laghrissi A, Juodėnas M, Tamulevičius T, et al. Magnetic-assisted sequential templated self-assembly of hybrid colloid nanoparticle systems. Nanoscale 2024;16:22167–77.

49. Fehse S, Das A, and Belder D. Integration of a recyclable silver substrate for in situ surface-enhanced Raman spectroscopy in digital microfluidics. Chemical Communications 2024;60:8252–5.

50. Liu D, Yang Z, Zhang L, et al. Cell-free biology using remote-controlled digital microfluidics for individual droplet control. RSC Advances 2020;10:26972–81.

51. Lobato IM and O’Sullivan CK. Recombinase polymerase amplification: basics, applications and recent advances. Trac Trends in Analytical Chemistry 2018;98:19–35.

52. Daher RK, Stewart G, Boissinot M, et al. Recombinase polymerase amplification for diagnostic applications. Clinical Chemistry 2016;62:947–58.

53. Li J, Macdonald J, and Von Stetten F. A comprehensive summary of a decade development of the recombinase polymerase amplification. Analyst 2019;144:31–67.

54. Piepenburg O, Williams CH, Stemple DL, et al. DNA detection using recombination proteins. PLoS Biology 2006;4:e204.

55. Shen HH, Fan SK, Kim CJ, et al. EWOD microfluidic systems for biomedical applications. Microfluidics and Nanofluidics 2014;16:965–87.

56. Munawar MA. Critical insight into recombinase polymerase amplification technology. Expert Review of Molecular Diagnostics 2022;22:725–37.

57. Kestin J, Sokolov M, and Wakeham WA. Viscosity of liquid water in the range-8 C to 150 C. Journal of Physical and Chemical Reference Data 1978;7:941–8.

58. Tian J, Zhang Y, Zhu L, et al. dsDNA/ssDNA-switchable isothermal colorimetric biosensor based on a universal primer and *ω* exonuclease. Sensors and Actuators B: Chemical 2020;323:128674.

59. Weiss B. Endonuclease II of Escherichia coli is exonuclease III. Journal of Biological Chemistry 1976;251:1896–901.

60. Zhang Y and Tanner NA. Isothermal amplification of long, discrete DNA fragments facilitated by single-stranded binding protein. Scientific Reports 2017;7:8497.

