## Supplementary Methods and Results for "Orchestrating Self-Replication in Artificial Cells through Digital Microfluidics"

### S1 Supplementary Movies

#### Movie S1: DMF-based replicator

Movie S1 shows cyclic droplet replication from Cycle 0 to Cycle 8 on the OpenDrop platform. The initial cycle (Cycle 0) begins with a methylene blue droplet (1 mM, ultra-pure water, 1% v/v silicone oil AR5). In each cycle, a tartrazine droplet (1 mM, ultra-pure water, 1% v/v silicone oil AR5) is added to dilute the droplet from the previous cycle. Droplets mix in a 2×2 electrode array and are then split along a predefined electrode path to achieve consistent serial dilution over 8 rounds. Note: tartrazine and methylene blue are assigned to opposite reservoirs compared to Fig. 2a.

### S2 Supplementary Materials and Methods

#### S2.1 Materials

Device fabrication required additional materials beyond the core OpenDrop components: 3M PET double-sided acrylic adhesive and 3M 9719 conductive adhesive transfer tape (3M, USA), 7 µm PVC film (Bacofoil, UK), Hylosil Instant Gasket Silicone Sealant (RS Components Ltd., UK), and PDMS Sylgard 184 Silicone Elastomer Kit containing base elastomer and curing agent (Dow Corning, USA). Surface treatment utilized ethanol (pure $\geq$ 99.8%, GC, Merck Life Science Limited, UK) and aluminum foil for contamination prevention (Thermo Fisher Scientific, UK). Enhanced RPA analysis required EDTA tetrasodium salt hydrate (98%, Scientific Laboratory Supplies Ltd., UK), DNA UltraPure Agarose, SYBR gold nucleic acid gel stain (Thermo Fisher Scientific, UK), Quick-Load 1 kb Plus DNA Ladder (New England BioLabs, UK), and custom primers synthesized by Integrated DNA Technologies, Belgium.

#### S2.2 OpenDrop Modifications

**1. Dielectric Film.** OpenDrop recommends an ETFE Film with a thickness of 12 µm. The Young-Lipmann's equation predicts that a higher dielectric constant ($\varepsilon$) and a smaller thickness result in a higher capacitance, which lowers the actuation voltage and improves the sensitivity of droplet movement. Therefore, we initially selected PTFE film ($\varepsilon \approx 2.1$) with a thickness of 5 µm, which has a dielectric constant similar to ETFE film ($\varepsilon \approx 2.6$). However, the PTFE film was prone wrinkling on application and would breakdown under high voltage (Fig. S1a). We ultimately selected a 7 µm PVC film ($\varepsilon \approx 3.1$), which provided a smoother surface, better plasticity, and sufficient dielectric strength, making it more suitable for reliable droplet manipulation (Fig. S1b).

**2. Droplet Channel.** A double-sided acrylic adhesive spacer with thickness of 278 µm was incorporated to support the top ITO glass and the bottom electrode arrays (Fig. S2a). The custom design is shown in Fig. S2b, where the double-sided acrylic adhesive spacer was cut to a frame shape. However, using only the frame spacer was insufficient to keep the top ITO glass flat during the sealing process, leading to variations in droplet volume across the array (outlined in red, Fig. S2c). Based on this custom design, we further applied the adhesive spacer to the central region of the non-electrode area (Fig. S2d). The improvement ensured that the top ITO glass remained flat, resulting in uniform droplet volumes across the array (Fig. S2e).

**3. Droplet Surface Tension.** In our experiments, silicone oil (5 cSt) is added to reduce the surface tension of the water-based solution, facilitating smooth injection into the reservoirs and sensitive droplet movement. We investigated the fusion and fission of silicone oil AR5 (5 cSt) in a methylene blue droplet (0.1 mM in ultra-pure water) at volume concentrations of 0%, 0.08%, 0.10%, 0.12%,

0.15%, 0.18%, and 0.20%, with droplet movement speeds ranging from 1 frame/s to 4 frames/s. The results indicated that 0.10% was the optimal concentration for the droplet. Different variations of the OpenDrop device were used in subsequent studies and the volume concentration of silicone oil AR5 (5 cSt) was adjusted to 1% v/v.

**4. Enclosed Chip.** A silicone gasket was applied to effectively seal the gaps between the frame and the electrode board.

**5. Medium Oil Selection.** Typical isothermal amplification on digital microfluidic systems reported to use silicone oil with a viscosity of 5 cSt as the medium oil. Here, we compared the silicone oil (5 cSt) and octamethyltrisiloxane with a lower viscosity of 1.2 cSt. Two methylene blue droplets (1 mM, ultra-pure water, 1% v/v silicone oil AR5) were split within the oils, and their behaviour was recorded. After droplets split in silicone oil (5 cSt), the daughter droplets were uneven, with a satellite droplet forming. In contrast, droplets splitting in octamethyltrisiloxane took advantage of the equal splitting of droplets.

**7. Temperature Calibration.** We compared the actual temperature with the preset temperature of the chip. An ultra-pure water droplet was placed on the heating region of the electrode board covered with a fluoropel-coated PVC film, where the actual temperature of the droplet was measured using an electronic thermometer (Fig. S3).

##### S2.2.1 Fabrication Process

First, the masking tape was applied on the bare electrode board to protect the circuit (Fig. S4a). A degassed PDMS premix (10:1 ratio of reagents A and B) was spread evenly on the board (Fig. S4b), covered with a PVC film and laminated (Fig. S4c). The assembly was cured at room temperature for 12 hours, with aluminium foil covering the surface to prevent dust contamination. Next, the drilled ITO glass was cleaned with ethanol and Milli-Q water, dried under nitrogen, and its conductive side was verified using a digital multimeter. Both the ITO glass with a conductive side and the electrode board with its PVC film facing upward were spin-coated with 60 µl FluoroPel PFC1601V at 2500 rpm for 30 seconds (Fig. S4d-e), followed by curing at 180 °C and 90 °C, respectively, for 20 minutes. A thickness of 278 µm 3M PET double-sided acrylic adhesive was applied to bond the electrode board and frame (Fig. S4f), followed by the gasket silicone sealant sealing the gaps between the PVC film and the frame (Fig. S4g). The sealant was cured at room temperature for 12 hours. Finally, one side of the conductive tape was bonded to the contact electrode P1 on the electrode board, while the other was attached to the conductive side of the ITO glass. The ITO glass was aligned with its coated surface facing downward toward the electrode board, carefully positioned at the centre of the frame (Fig. S4h).

#### S2.3 Optimization of RPA for DMF Applications

For optimized DNA amplification in the DMF environment, we investigated the effect of enzyme concentration on reaction kinetics. For a standard reaction, reservoir A was filled with a RPA reaction mixture (1.68 µM forward primer, 1.68 µM reverse primer, 0.48 µM probe, 28.23 mM Tris-HCl, 113.2 µM dNTPs, 1.7 mM ATP, 56.6 mM potassium acetate, 1.13 mM DTT, 28.23 mM phosphocreatine, and 0.1% Tween 20 (v/v), pH 7.9; one lyophilized enzyme pellet). Reservoir B was filled with $Mg(OAc)_2$ solution (56 mM magnesium acetate, 67.76 mM Tris-HCl, 271.04 µM dNTPs, 4.07 mM ATP, 135.52 mM potassium acetate, 2.71 mM DTT, 67.76 mM phosphocreatine, and 0.1% Tween 20 (v/v), pH 7.9). Reservoir C was filled with 0.32 nM solution of positive control DNA template. The effect of enzyme concentration on DNA amplification rate was investigated by doubling the enzyme content in Reservoir A (two lyophilized enzyme pellets instead of one), while maintaining all other components unchanged.

For the experiment described in Fig. 4 reservoir A was filled with a RPA reaction mixture (1.68 µM forward primer, 1.68 µM reverse primer, 0.48 µM probe, 28.23 mM Tris-HCl, 113.2 µM dNTPs, 1.7 mM ATP, 56.6 mM potassium acetate, 1.13 mM DTT, 28.23 mM phosphocreatine, and 0.1% Tween 20 (v/v), pH 7.9; lyophilized enzyme). Reservoir B was filled with $Mg(OAc)_2$ solution (56 mM magnesium acetate, 67.76 mM Tris-HCl, 271.04 µM dNTPs, 4.07 mM ATP, 135.52 mM potassium acetate, 2.71 mM DTT, 67.76 mM phosphocreatine, and 0.1% Tween 20 (v/v), pH 7.9) or EDTA solution (160 mM EDTA, 67.76 mM Tris-HCl, 271.04 µM dNTPs, 4.07 mM ATP, 135.52 mM potassium acetate, 2.71 mM DTT, 67.76 mM phosphocreatine, and 0.1% Tween 20 (v/v), pH 7.9). Reservoir C was filled with 0.32 nM solution of positive control DNA template.

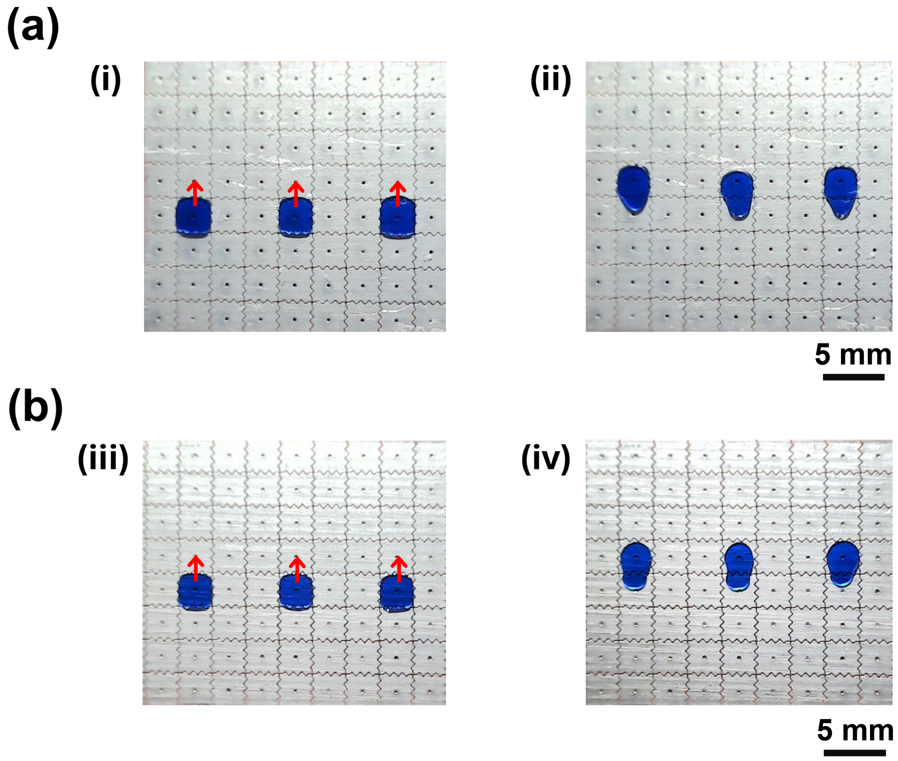


Figure S1: Comparison of methylene blue droplet (10 mM, ultra-pure water, 1% v/v silicone oil AR5) actuation on chips with different dielectric films. (a) Droplet actuation on a chip featuring a 5 µm PTFE dielectric film forms wrinkles on the board, and (b) Droplet actuation on a chip with a wrinkle-free 7 µm PVC dielectric film.

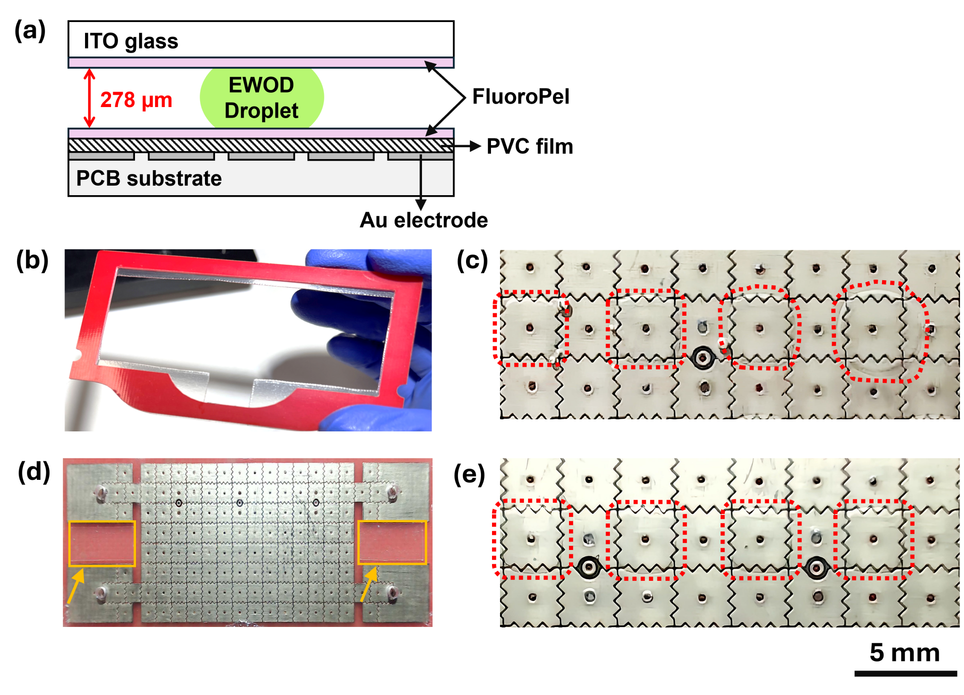


Figure S2: Optimization of droplet channel within the chip. (a) Schematic cross-sectional view of the digital microfluidic chip. An ITO glass slide is coated with a FluoroPel polymer layer, a 278 μm spacer provides for droplet movement, a FluoroPel-coated PVC dielectric film is mounted on the Au electrode array and PCB substrate. (b) Custom design contained a double-sided acrylic adhesive spacer, which was cut into the shape and attached to the frame to support the top ITO glass and the bottom electrode board. (c) Photograph of variations in ultra-pure water droplet volumes (red dotted line) due to ITO glass tilting in the absence of additional adhesive in the non-electrode region. (d) Optimized design incorporated extra double-sided adhesive (indicated by the orange rectangle and arrow) in the central non-electrode region to maintain the flatness of the top ITO glass and prevent tilting. (e) Photograph of uniform ultra-pure water droplet volumes (red dotted line) after applying the optimized configuration.

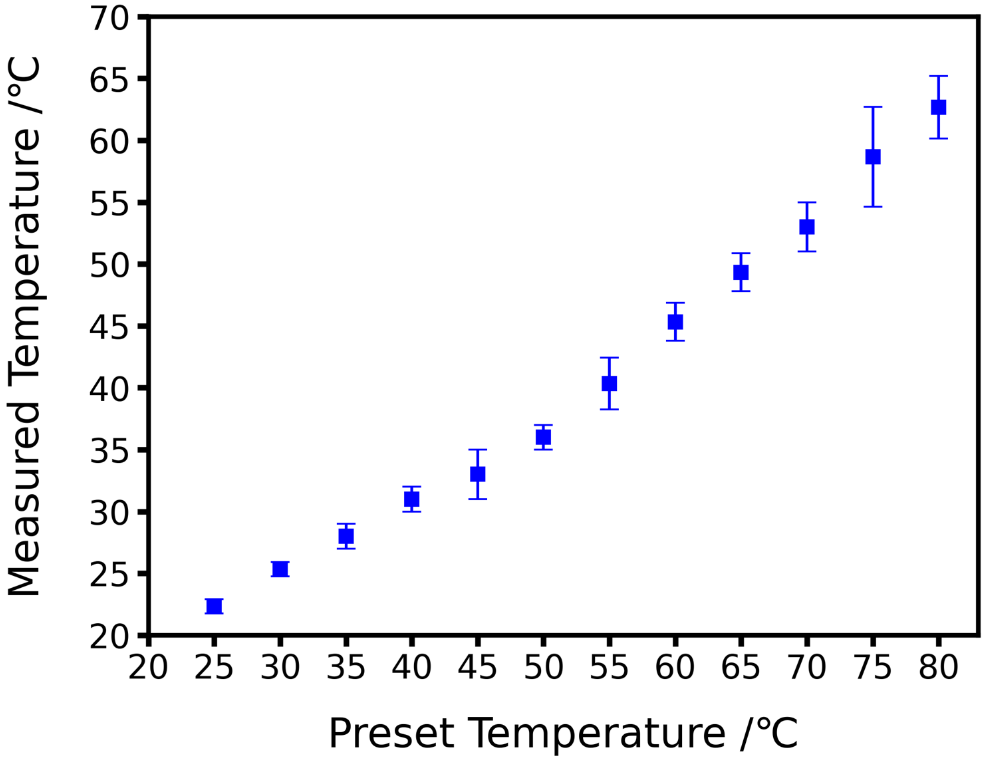


Figure S3: Calibration of the chip heater by comparing the preset and measured temperatures ranging from 25 to 80 °C. Preset values were set via software, and actual droplet temperatures were measured using an electronic thermometer. Error bars represent the standard deviation from three replicates.

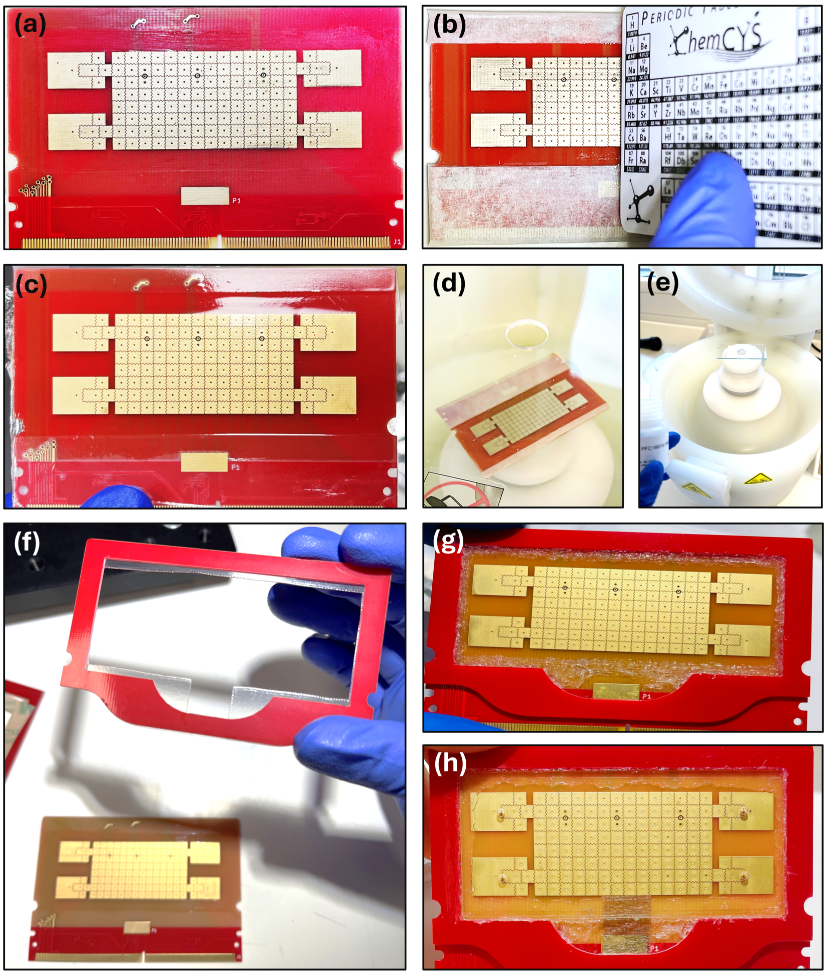


Figure S4: Photographs of OpenDrop chip fabrication process. (a) The bare electrode board is covered with masking tape for protection. (b) A PDMS premix is evenly spread over the electrode board. (c) The PDMS-coated electrode board was laminated using PVC film and a preheated laminator. (d) A hydrophobic layer of Fluoropel PFC1601V is applied via spin-coating onto the electrode board with the flat dielectric PVC film. (e) A Fluoropel layer is also applied via spin-coating on the conductive side of ITO glass. (f) A double-sided acrylic adhesive of thickness 278 µm was used to seal the frame and the Fluoropel-coated electrode board. (g) A gasket silicone sealant was applied to seal the gap between the frame and the electrode board. (h) The final OpenDrop chip with the ITO glass positioned in the centre of the frame.

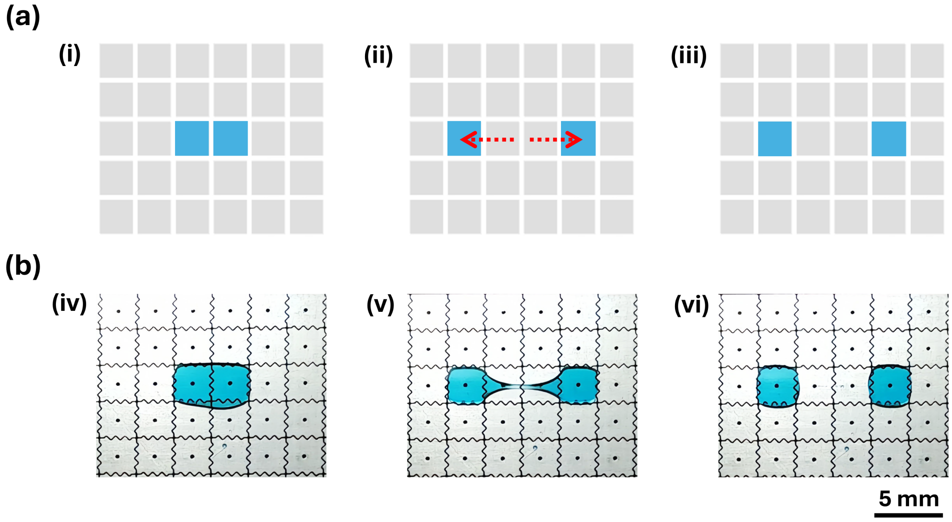


Figure S5: Droplet division via electrode activation. (a) Schematic illustrating droplet splitting via electrode activation: (i) A droplet is positioned across two adjacent electrodes; (ii) Activation of the outer electrodes while deactivating the positioned electrodes; (iii) The droplet divides into two equal daughter droplets. (b) Corresponding photographs showing stages (iv), (v), and (vi), which respectively match stages (i), (ii), and (iii) from the schematic. The droplet contains 1 mM methylene blue in ultrapure water with 1% v/v silicone oil AR5. Scale bar: 5 mm.

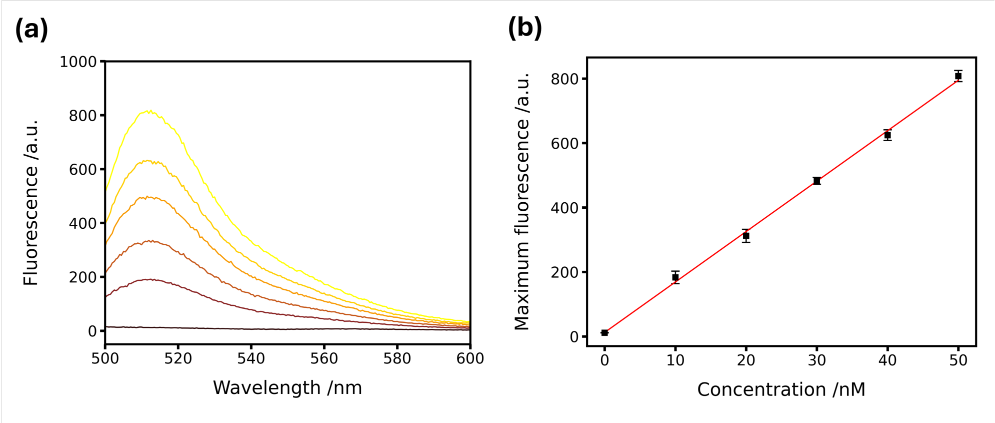


Figure S6: Calibration for standard fluorescein solutions. (a) Fluorescence spectra of fluorescein solutions at concentrations ranging from 0 nM to 50 nM (0 nM: black; 10 nM to 50 nM: red to yellow gradient). Fluorescence measurements were performed using a fluorometer (Cary Eclipse, excitation wavelength 490 nm, emission wavelength 513 nm, excitation slit 5 nm, emission slit 5 nm, PMT voltage 1000 V). (b) A linear calibration curve was generated by plotting the maximum fluorescence intensity against fluorescein concentration. The fitted equation is $y = 15.6467x + 12.4396$ ($R^2 = 0.9970$), where y represents the maximum fluorescence intensity and x represents the fluorescein concentration (nM).

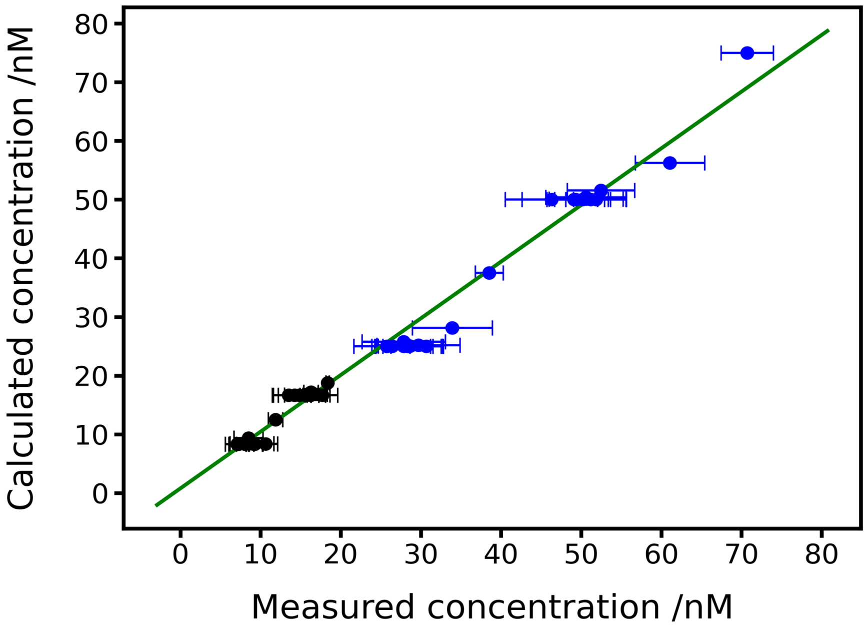


Figure S7: Relationship between measured and calculated fluorescein concentrations in cyclic replication. Black data points represent 25 nM fluorescein diluted with buffer, and blue data points represent 75 nM fluorescein with buffer. Linear regression analysis ($y = 0.9656x + 0.7908$ with $R^2 = 0.9816$) demonstrates excellent agreement between experimental and theoretical values.

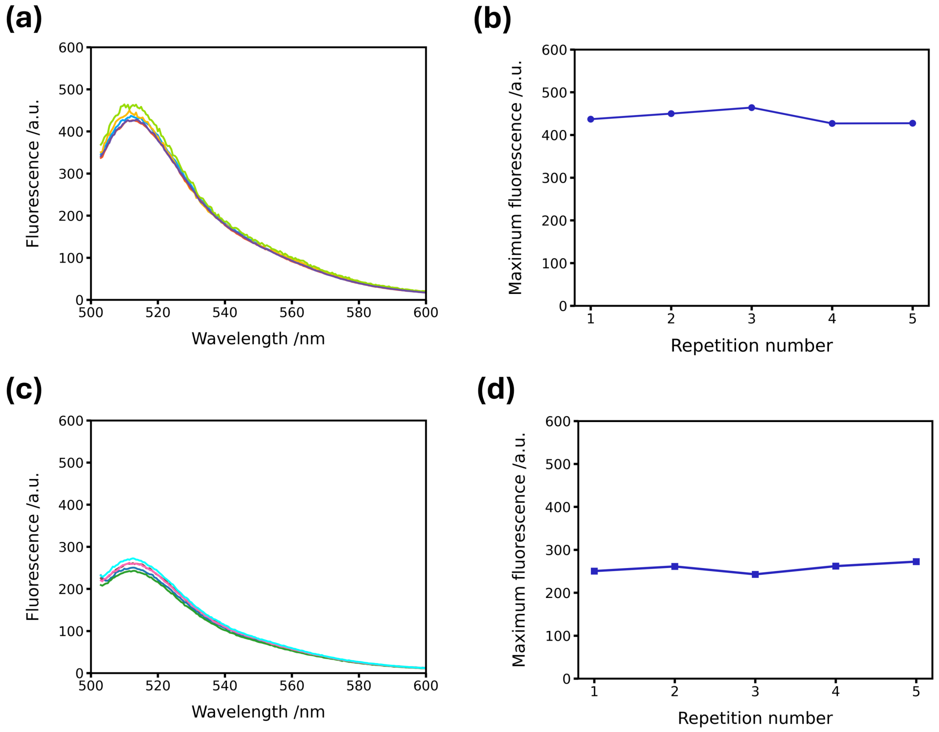


Figure S8: Reproducible mixing and division of individual droplets. A fluorescein solution from reservoir A (25 nM in 0.1 M Tris-HCl buffer, pH 8.0, 1% (v/v) silicone oil AR5) was repeatedly mixed with buffer solution from reservoir B (0.1 M Tris-HCl, pH 8.0, 1% (v/v) silicone oil AR5). Fluorescence spectra of individual droplets over five repeated cycles for (a) mixing and (c) mixing-division. Corresponding plots of maximum fluorescence intensity versus repetition number are shown in (b) for mixing and (d) for mixing-division. The coefficient of variation (CV) was 3.72% (mean $\pm$ SD: 27.39 $\pm$ 1.02 nM) for mixing and 4.65% (mean $\pm$ SD: 15.68 $\pm$ 0.73 nM) for mixing-division, indicating high reproducibility (CV $<$ 5%).

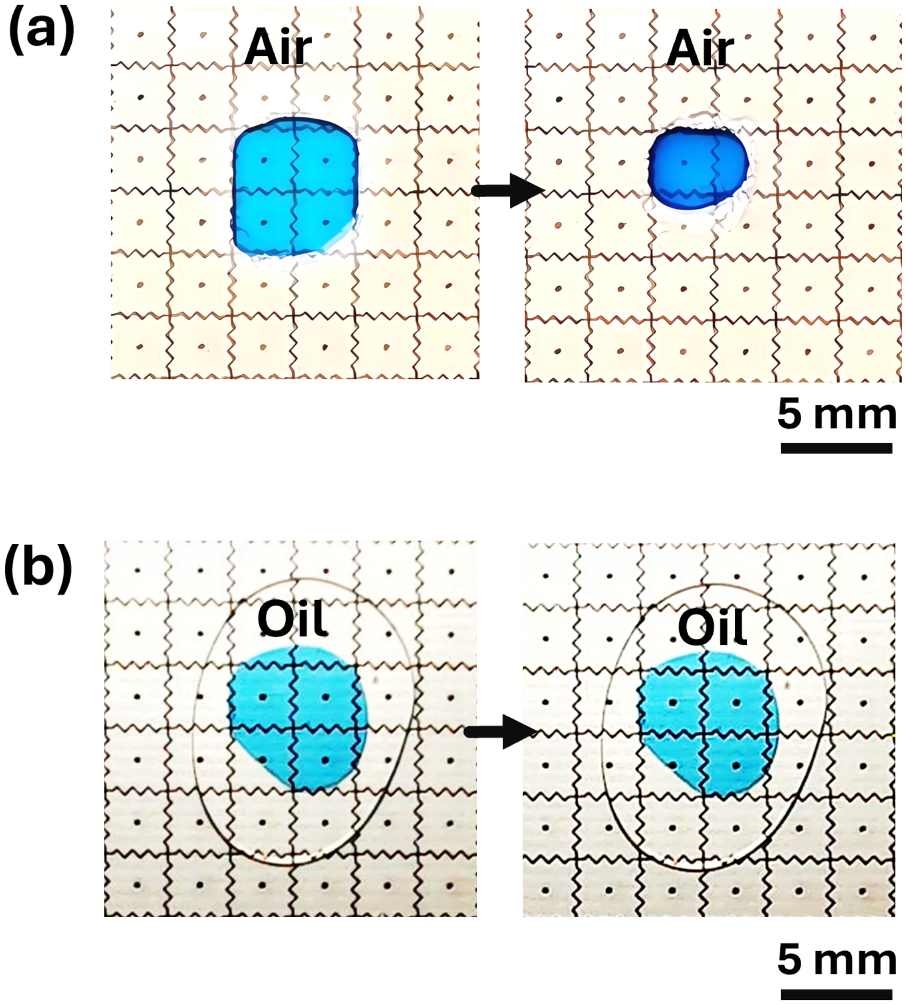


Figure S9: Photographs comparing droplet evaporation in air and oil phase. Methylene blue droplets (1 mM, ultrapure water, 1% v/v silicone oil AR5) were heated at 40 °C under two conditions: (a) exposed to air and (b) immersed in an oil phase. Images were taken at 0 minutes (left) and 60 minutes (right). Scale bar: 5 mm.
